# DNA Polymerase β Accelerates Cellular DNA Base Excision Repair by Suppressing Excessive PARP1 Engagement

**DOI:** 10.64898/2026.07.20.739154

**Authors:** Annie A. Demin, Marek Adamowicz, Jan Brazina, Amit Gautam, Keith W Caldecott

## Abstract

DNA polymerase beta (POLβ) is required for rapid rates of cellular DNA base excision repair (BER). However, the reason for this requirement is unclear, because other DNA polymerases can replace POLβ, in vitro. Here, we have identified the essential role of POLβ during cellular BER. As expected, POLβ deletion in human RPE-1 cells resulted in the rapid accumulation of DNA strand break intermediates during incubation with the monofunctional alkylating agent, methyl methanesulphonate (MMS). However, this accumulation was not detected in cells that also lack PARP1, indicating that POLβ is required for BER only if PARP1 is present. This result is reminiscent of the essential role of XRCC1 during BER, which is to suppress the excessive engagement and activity of PARP1 at BER intermediates and thereby enable their access and repair by other enzymes. Indeed, we found that POLβ is required to prevent excessive PARP1 engagement and activity during BER, and that XRCC1 and POLβ fulfil this function together. Finally, similar to XRCC1, loss of POLβ leads to persistent transcriptional suppression during MMS-induced BER, and this suppression is alleviated by treatment with PARP inhibitor. In summary, we show here that the essential role of POLβ during cellular BER is to suppress excessive PARP1 engagement and activity, and thereby maintain rapid rates of this important DNA repair process.

## Introduction

DNA base excision repair (BER) is an evolutionary conserved process that replaces missing or damaged nucleobases with their intact undamaged counterpart (1–3). Minimally, this process involves removal of damaged bases by a DNA glycosylase, incision of abasic sites by an AP endonuclease, DNA gap filling by a DNA polymerase, and DNA ligation by a DNA ligase. In mammalian cells, a number of accessory factors are present that increase the rate of this process, such as PARP1/PARP2(4–6) and the scaffold protein, XRCC1 (7–9). In mammalian cells, the DNA polymerase primarily involved in BER is DNA polymerase β (POLβ). This was first suggested by biochemical experiments in which permeabilised nuclei were treated with different genotoxins, and DNA repair synthesis was measured in the presence of different DNA polymerase inhibitors (10–13). Subsequently, a role for POLβ in BER was suggested by experiments in which BER reactions were reconstituted in vitro using cell extracts (14–21) or purified proteins (22, 23). Most importantly, reduced rates of BER are readily detected in living cells in which POLβ is deleted (24–26).

POLβ interacts with the scaffold protein XRCC1 (22, 27, 28), assembles into BER multiprotein complexes (29, 30), and possesses two catalytic activities, each of which can facilitate a key step of BER. These are a 5’-deoxyribose phosphate lyase (dRPase) activity that removes the 5’-terminal sugar-phosphate created by APE1, and a nucleotide transferase activity that conducts DNA gap filling (23, 31). Of these two activities, it is the loss of dRPase activity that has the greatest impact on cellular BER (21). Despite its clear utility, however, it remains unclear why POLβ is required for BER, because both the DNA gap filling and dRPase activities are present in other DNA polymerases, such as POLλ and POLι (32, 33). Indeed, either of these enzymes can substitute for POLβ in BER reactions employing purified proteins, in vitro (32, 33). In addition, mammalian cells possess a long-patch sub-pathway of BER that can function independently of POLβ, by employing POLδ and FEN1 endonuclease to conduct gap filling and to displace and excise 5’-deoxyribose termini prior to the final step of DNA ligation (17, 18, 34, 35). Moreover, long-patch BER operates in human cells more frequently than previously thought, and can bypass the requirement for POLβ during BER, but nonetheless cannot support the rapid initial rate of BER that is facilitated by POLβ (26, 36–38). Consequently, here, we have addressed directly why POLβ is required for rapid rates of BER in human cells.

## Results

We showed recently that XRCC1 accelerates BER primarily by suppressing excessive PARP1 engagement at BER intermediates, which can impede the recruitment of other BER proteins, including XRCC1 itself (39). As a result of this, XRCC1 recruitment into chromatin is greater in *PARP1^−/−^* RPE-1 cells than in wild type RPE-1 cells, during MMS treatment (39). While this observation identified the primary mechanism by which XRCC1 accelerates BER, it questioned the notion that PARP activity is required for XRCC1 recruitment, during BER. To address this, we compared the MMS-induced accumulation of XRCC1 into chromatin in wild type RPE-1 cells and in RPE-1 cells lacking either PARP1 and/or PARP2. Whereas deletion of PARP1, as expected, increased XRCC1 accumulation during MMS treatment, deletion of PARP2 did not have a significant effect (Fig.1A). More importantly, the deletion of both PARP1 and PARP2 greatly reduced XRCC1 accumulation (Fig.1A). Thus, either PARP1 or PARP2 activity is required for the recruitment of XRCC1 during BER, which subsequently then limits the extent of PARP1 engagement at BER intermediates.

**Figure 1.**
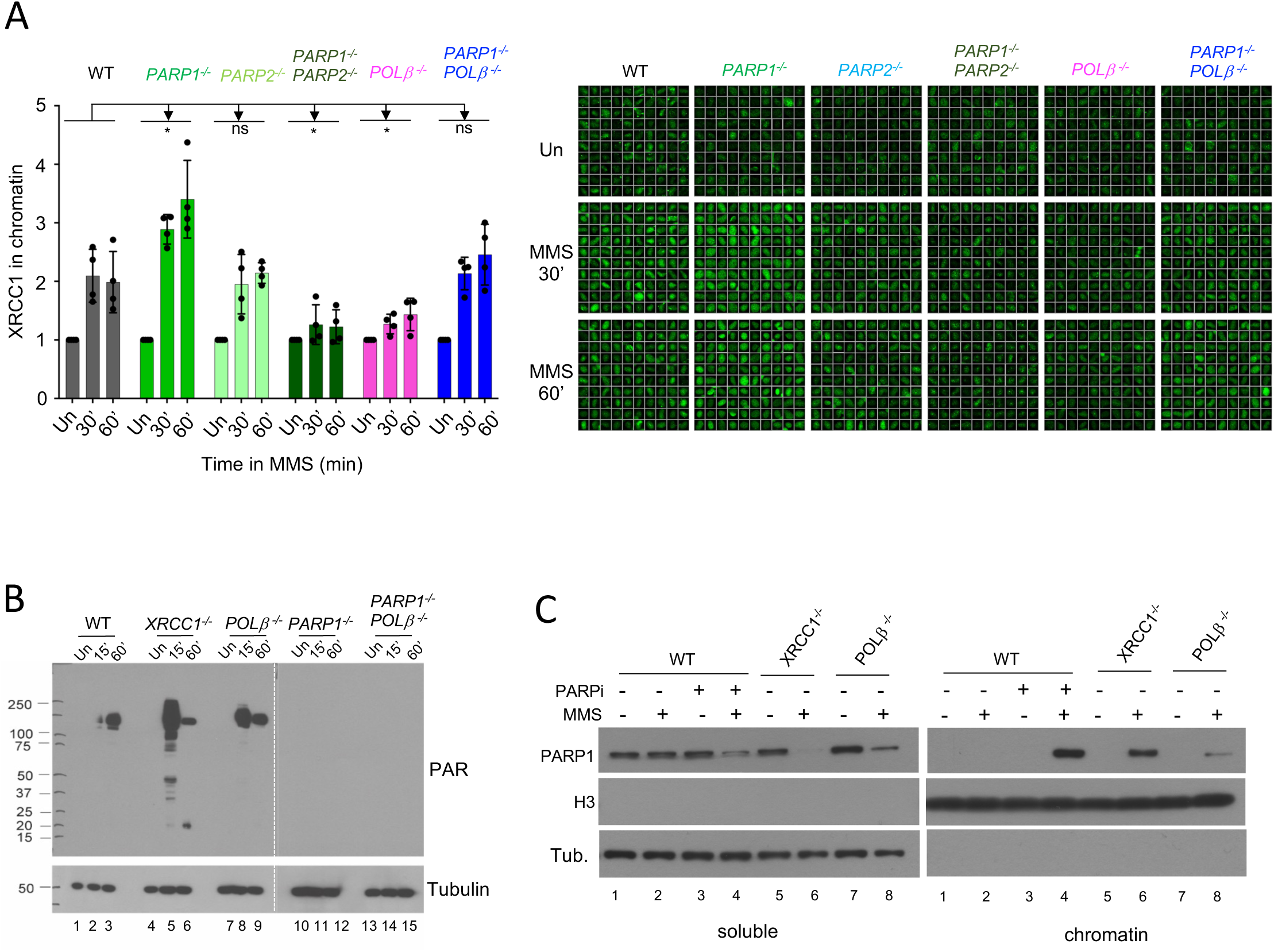
PARP1 chromatin retention and hyperactivation during BER in *POL*β *^−/−^* cells. **(A),** The presence of XRCC1 protein in chromatin, as defined by its resistance to extraction with detergent, was detected by indirect immunofluorescence in the indicated wild type, *PARP1^−/−^*, *PARP2^−/−^*, *PARP1^−/−^/PARP2^−/−^*, *POL*β *^−/−^* (clone #AB1), and *PARP1^−/−^*/*POL*β *^−/−^* (clone #1) mutant RPE-1 cell lines following the indicated incubation periods in 0.1 mg/ml MMS. The amount of detergent-insoluble XRCC1 was quantified, relative to that in untreated cells, by automated high-content scanR imaging. Data on the left are the mean (+/−SD) of four independent experiments. The panel on the right is a gallery of representative scanR images of XRCC1 immunostaining in chromatin, with each large square containing 100 individual cells. **(B)**, PARP1 auto-ribosylation in wild type, *XRCC1^−/−^*, *POL*β *^−/−^* (clone #AB1), *PARP1^−/−^*, and *PARP1^−/−^/ POL*β *^−/−^* (clone #1) RPE-1 cells. RPE-1 cells during treatment with 0.1 mg/ml MMS, detected by the poly (ADP-ribose)-specific detection reagent MABE1031. Tubulin was immunostained for a loading control. **(C)**, PARP1 levels in cell-equivalent aliquots of detergent-soluble and detergent-resistant (chromatin-containing) fractions of the indicated wild type (WT), *XRCC1^−/−^*, and *POL*β *^−/−^* (clone #AB1) mutant RPE-1 cell lines, measured by western blotting. Histone H3 (H3) and Tubulin were immunostained for loading controls. Cells were incubated or not with 10 μM PARP inhibitor (KU0058948) and/or MMS (0.1 mg/ml) for 1 h, as indicated, prior to subcellular fractionation. Representative blots are shown.

We noticed in these experiments that the recruitment of XRCC1 into chromatin during MMS-induced BER was also reduced in *POL*β *^−/−^* cells, but occurred at wild type levels in *POL*β *^−/−^* cells in which PARP1 was also deleted (Fig.1A). This suggested that, similar to XRCC1, POLβ might be required to supress excessive PARP1 engagement during BER. In XRCC1-defective cells, excessive PARP1 engagement during MMS-induced BER is accompanied by a rapid burst of PARP1 auto-ribosylation, followed by a subsequent decline in PARP1 activity and the “trapping” of PARP1 in chromatin as a result of NAD^+^ depletion (39). We therefore examined whether this was also true in *POL*β *^−/−^* cells. Indeed, similar to *XRCC1^−/−^* cells, *POL*β *^−/−^* RPE-1 cells exhibited much higher levels of ADP ribosylation than did wild type cells at early times (15 min) during MMS treatment (Fig.1B, compare lanes 2, 5 & 8). This elevated ADP-ribosylation activity in *POL*β *^−/−^* cells was the result of PARP1, because it was absent from cells in which PARP1 was deleted (Fig.1B, lanes 13-15). Moreover, elevated MMS-induced ADP-ribosylation in *POL*β *^−/−^* cells was accompanied by an eventual accumulation of PARP1 in chromatin, albeit not to the same extent as that observed in *XRCC1^−/−^* cells or in wild type cells treated with PARP inhibitor (Fig.1C, compare lanes 2, 4, 6, & 8). Thus, similar to XRCC1, POLβ is required to supress PARP1 hyperactivation and ‘trapping’ during MMS-induced BER.

To examine the importance of POLβ for protecting BER from excessive engagement and “trapping” of PARP1, we measured the accumulation of BER intermediates during MMS-induced BER using alkaline comet assays. In agreement with our previous work (39), DNA strand breaks rapidly accumulated in *XRCC1^−/−^* cells to much higher levels than in wild type cells during MMS treatment, and this increased accumulation was ablated by deletion of PARP1 (Fig.2A, compare WT, *XRCC1^−/−^*, & *XRCC1^−/−^*/*PARP1^−/−^* cells). The DNA strand breaks detected here reflected BER intermediates because their appearance was prevented by deletion of methyl purine/alkyl adenine DNA glycosylase (MPG/AAG), the DNA glycosylase responsible for initiating excision of the methylated DNA bases (Fig.2B). Importantly, *POL*β *^−/−^*cells also exhibited a much greater increase in MMS-induced DNA breaks than did wild type cells, albeit to a lesser extent than *XRCC1^−/−^* cells (Fig.2A, compare WT, *XRCC1^−/−^*, & *POL*β *^−/−^*cells). Moreover, this increase was similarly suppressed by deletion of PARP1 (Fig.2A, *POL*β*^−/−^*/*PARP1^−/−^* cells). This result was not an artifact of gene editing or clonal selection, because we observed similar results in two independent *POL*β *^−/−^* clones and by employing PARP1 siRNA (Fig.3A). These data indicate that, similar to XRCC1, POLβ suppresses excessive PARP1 engagement during BER, which otherwise impedes DNA repair and leads to the accumulation of SSB intermediates.

**Figure 2.**
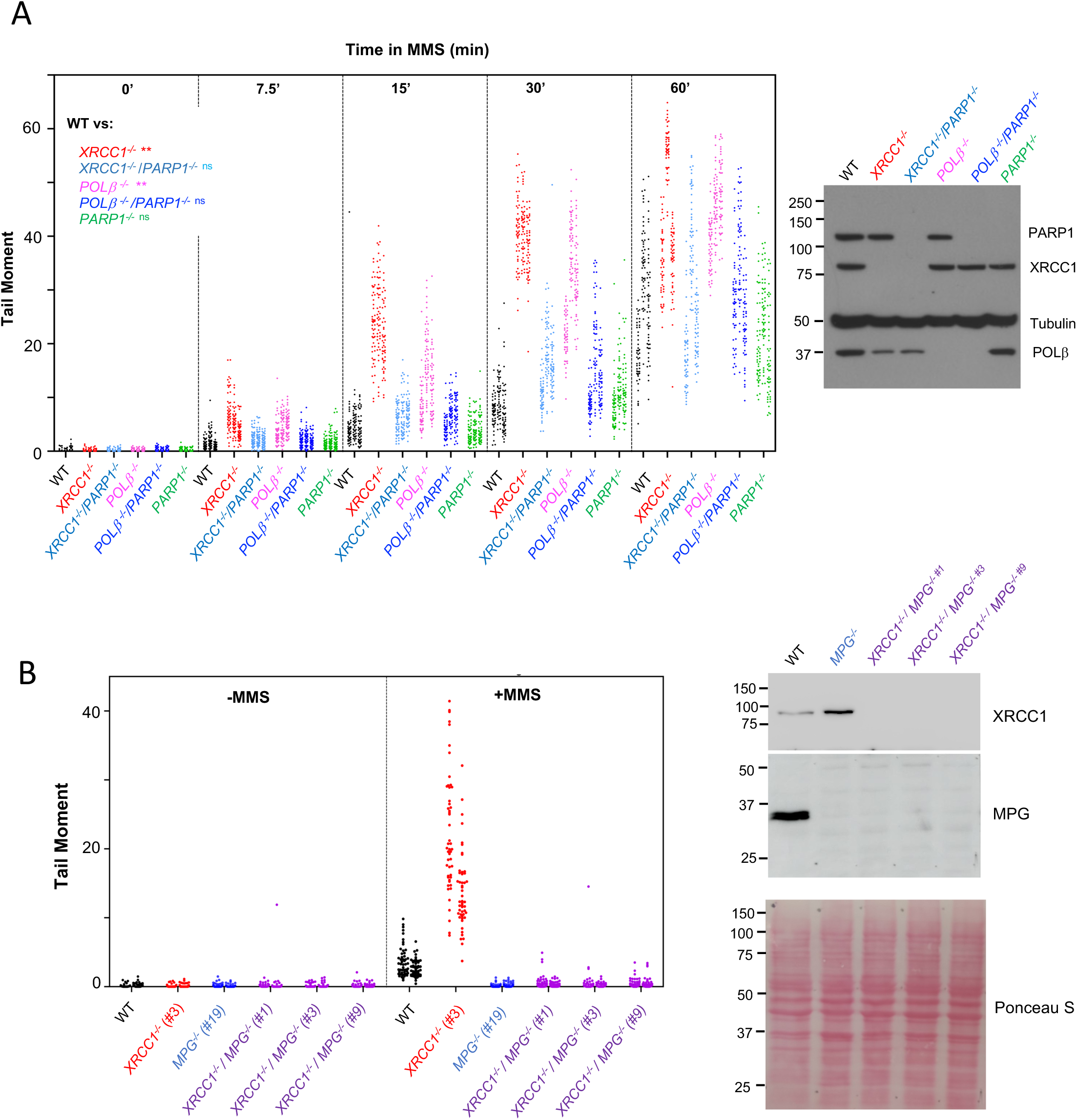
PARP1 is responsible for the elevated accumulation of DNA strand breaks during BER in *POL*β *^−/−^* cells. **(A)**, *Left,* DNA strand breaks quantified by alkaline comet assays in wild type, *PARP1^−/−^*, *XRCC1^−/−^*, *XRCC1^−/−^/PARP1^−/−^*, *POL*β ^−/−^ (clone #AB1), and *PARP1^−/−^/ POL*β *^−/−^* (clone #1) RPE-1 cells at the indicated times during treatment with 0.1 mg/ml MMS. Plotted data are the individual comet tail moments (an arbitrary measure of DNA strand breakage) of fifty cells per sample per experiment, with the fifty tail moments for each experiment plotted vertically and three independent experiments plotted side by side. Statistically significant differences between wild type and the indicated mutant cell lines were determined by two-way ANOVA of the mean tail moments from the three independent experiments with Dunnett’s multiple comparisons test (ns, not significant; **p<0.01). *Right*, western blot showing levels of PARP1, XRCC1, POLβ, and tubulin loading control, in the indicated mutant cells. **(B)**, MMS-induced DNA breaks are intermediates of BER initiated by MPG. DNA strand breaks measured by alkaline comet assays in the indicated wild type (WT), *XRCC1^−/−^ (*clone #3), *MPG^−/−^* (clone #19), and *XRCC1^−/−^/MPG^−/−^* (clones #1, #3, & #9) cell lines in the absence of MMS treatment or 1 hr after treatment with 0.1 mg/ml MMS. Plotted data are the individual comet tail moments of fifty cells per sample per experiment, with the fifty tail moments for each experiment plotted vertically and two independent experiments plotted side by side. *Right*, western blot showing levels of XRCC1 and MPG/AAG in the indicated mutant cells. Ponceau S staining of the membrane was employed as a loading control.

**Figure 3.**
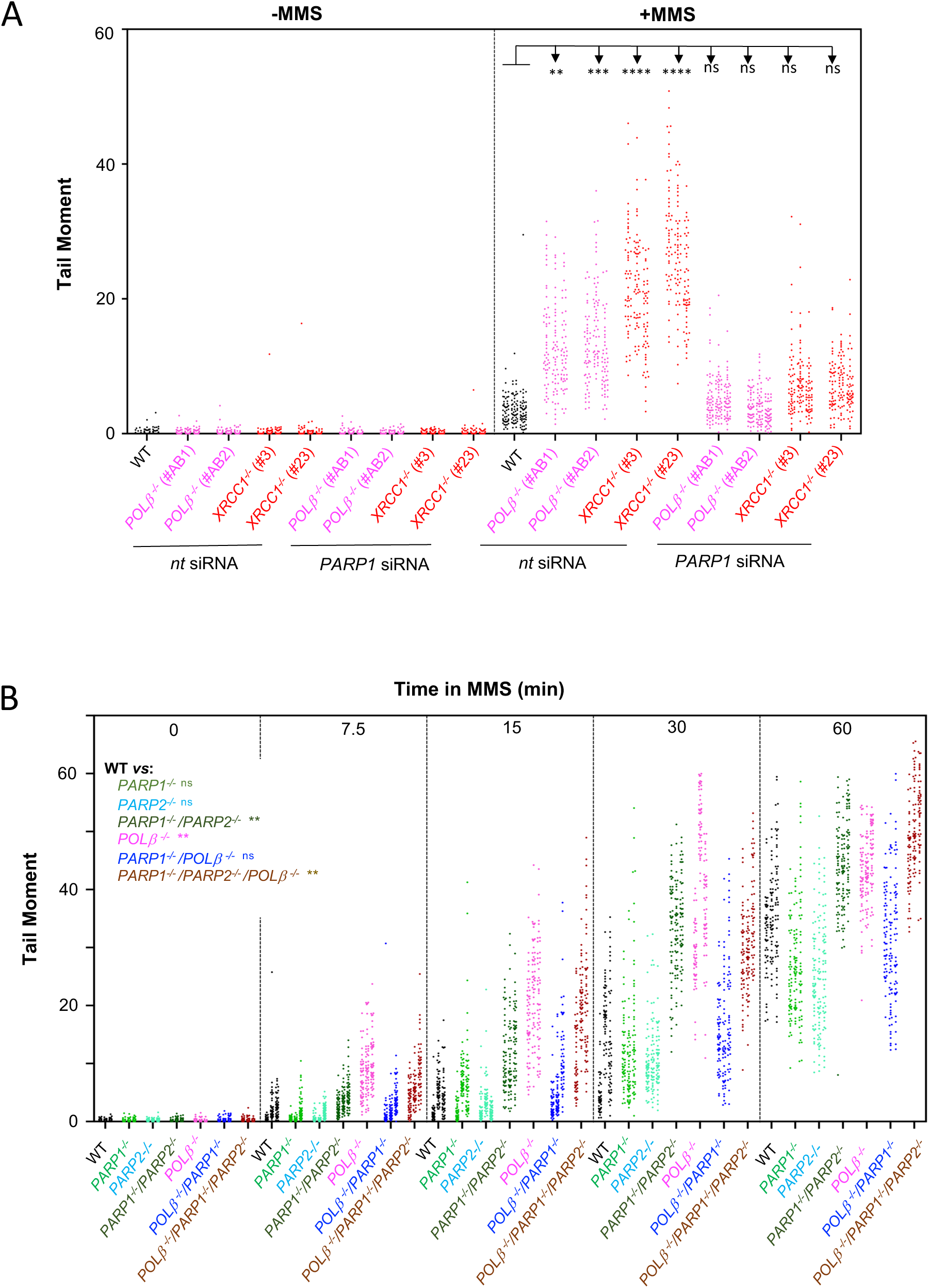
PARP2 can accelerate BER, in the absence of PARP1. **(A),** DNA strand breaks quantified by alkaline comet assays in wild type, *POL*β ^−/−^ (clones #AB1 & #AB2), and *XRCC1^−/−^* (clones #3 & #23) RPE-1 cells either without treatment (“un”) or 15 min after treatment with 0.1 mg/ml MMS. Cells were transfected with control non-targeting siRNA or PARP1 siRNA, prior to treatment, as indicated. Data plotted are the individual comet tail moments of fifty cells per sample per experiment, with the fifty tail moments for each experiment plotted vertically and three independent experiments plotted side by side. Statistically significant differences between wild type and the indicated mutant cell lines were determined by one-way ANOVA of the mean tail moments from the three independent experiments with Sidak’s multiple comparisons test (ns, not significant; **p<0.01; ***p<0.001; ****p<0.0001). **(B)**, DNA strand breaks quantified by alkaline comet assays in wild type, *PARP1^−/−^*, *PARP2^−/−^*, *PARP1^−/−^/PARP2^−/−^*, *POL*β ^−/−^ (clone #AB1), *POL*β *^−/−^*/*PARP1^−/−^* (clone #1), and *POL*β ^−/−^/*PARP1^−/−^/PARP2^−/−^* (clone #8) RPE-1 cells at the indicated times during treatment with 0.1 mg/ml MMS. Plotted data are the individual comet tail moments of fifty cells per sample per experiment, with the fifty tail moments for each experiment plotted vertically and three independent experiments plotted side by side. Statistically significant differences between wild type and the indicated mutant cell lines were determined by two-way ANOVA of the mean tail moments from the three independent experiments with Dunnett’s multiple comparisons test (ns, not significant; **p<0.01).

The above data indicate that, similar to XRCC1, the essential role of POLβ during BER is to suppress excessive PARP1 engagement and activity. It should be noted, however, that some PARP activity is required during BER, because while SSBs accumulated to wild type levels in *PARP1^−/−^* and *PARP2^−/−^* cells, they did so to higher levels in *PARP1^−/−^*/*PARP2^−/−^* cells (Fig.3B compare WT & *PARP1^−/−^*/*PARP2^−/−^* cells). Consistent with the above results, POLβ deletion did not increase the MMS-induced accumulation of DNA breaks during BER, in *PARP1^−/−^*/*PARP2^−/−^* cells (Fig.3B compare *PARP1^−/−^*/*PARP2^−/−^* & *POL*β *^−/−^/PARP1^−/−^*/*PARP2^−/−^*cells).

To examine whether XRCC1 and POLβ regulate PARP1 separately or together, we created cells lacking both proteins. As observed above, MMS-induced DNA strand breaks accumulated to higher levels than WT in both *XRCC1^−/−^* and *POL*β *^−/−^* cells, with the highest level detected in *XRCC1^−/−^* cells (Fig.4A). More importantly, DNA strand breaks accumulated in *XRCC1^−/−^* and *XRCC1^−/−^/POL*β *^−/−^* cells to a similar level, suggesting that XRCC1 and POLβ function together to suppress excessive PARP1 engagement during BER (Fig.4A).

**Figure 4.**
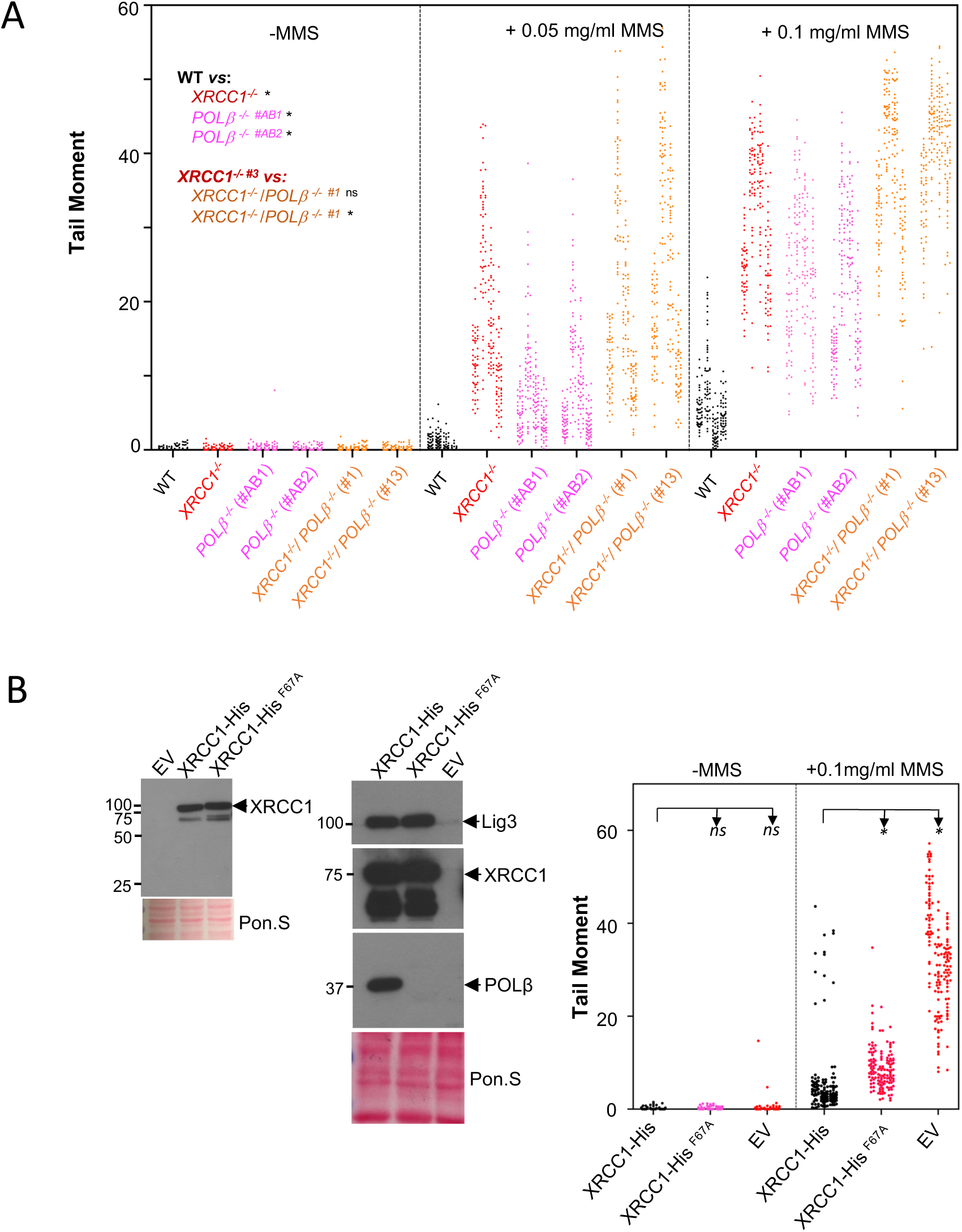
XRCC1 and POLβ function together to suppress excessive PARP1 engagement, during BER. **(A)**, DNA strand breaks quantified by alkaline comet assays in wild type, *XRCC1^−/−^*(clone #3), *POL*β ^−/−^ (clones #AB1 & #AB2), and *XRCC1^−/−^*/*POL*β ^−/−^ (clones #1 & #13) RPE-1 cells without treatment (“un”) or 15 min after treatment with the indicated concentrations of MMS. Data plotted are the individual comet tail moments of fifty cells per sample per experiment, with the fifty tail moments for each experiment plotted vertically and four independent experiments plotted side by side. Statistically significant differences between wild type and the indicated mutant cell lines were determined by two-way ANOVA of the mean tail moments from the three independent experiments with Dunnett’s multiple comparisons test (ns, not significant; *p<0.05). **(B)**, Mutation of the POLβ-binding domain in XRCC1 increases the accumulation of MMS-induced DNA breaks during BER. *Left panel*, western blots of total cell extract from wild type RPE-1 cells and *XRCC1^−/−^*(clone #23) RPE-1 cells stably transfected with either empty vector (“EV”) or plasmid encoding wild-type histidine-tagged human (XRCC1-His) or mutant histidine-tagged XRCC1 harbouring the mutation F67A (XRCC1-His^F67A^). *Middle panel*, western blots of anti-XRCC1 immunoprecipitates from the indicated cell lines. Note that XRCC1^F67A^ is unable to efficiently co-precipitate POLβ, but co-precipitates wild type levels of LIG3. *Right panel*, DNA strand breaks quantified in the indicated cell lines, either in the absence of MMS treatment or 15 min after treatment with 0.1 mg/ml MMS. Plotted data are the individual comet tail moments of fifty cells per sample per experiment, with the tail moments from each experiment plotted vertically and three independent experiments plotted side-by-side. Statistically significant differences between wild type and the indicated mutant cell lines were determined by two-way ANOVA of the mean tail moments from the three independent experiments with Dunnett’s multiple comparisons test (ns, not significant; *p<0.05).

Since XRCC1 and POLβ directly interact (22, 27, 40), we examined whether this interaction is required for the role of these proteins in suppressing excessive PARP1 engagement during MMS-induced BER. To do this, we employed *XRCC1^−/−^* RPE-1 cells stably expressing either wild type histidine-tagged XRCC1 (XRCC1-His) or mutant histidine-tagged XRCC1 (XRCC1-His^F67A^) harbouring an amino-terminal point mutation that disrupts the interaction with POLβ (41). As expected, affinity precipitation of wild-type XRCC1-His protein from human cell extracts co-recovered POLβ, whereas affinity precipitation of XRCC1-His^F67A^ did not (Fig.4B, left and middle panels). Importantly, although XRCC1-His^F67A^ reduced the accumulation of MMS-induced DNA breaks in *XRCC1^−/−^* cells, it did not restore these breaks to wild type levels, suggesting that the interaction of XRCC1 with POLβ is important for normal rates of BER (Fig.4B, right panel). The partial rescue of BER by XRCC1-His^F67A^ likely reflects that this protein retains the central BRCT domain, which can suppress PARP1 activity by direct binding to poly (ADP-ribose), and is required by XRCC1 to maintain rapid rates of BER (39, 42, 43). These data are consistent with previous results in CHO cells, in which mutation of the POLβ binding site in XRCC1 similarly reduced the rate of this process (44–46).

We showed recently that PARP1 hyperactivation in human *XRCC1^−/−^* cells during BER leads to an inability to maintain or restore transcriptional activity following MMS treatment (43). This phenomenon reflects aberrant recruitment of the ubiquitin protease USP3, triggered by elevated/persistent poly (ADP-ribose) synthesis at BER intermediates, which leads to deregulated histone de-ubiquitination and prolonged transcriptional inhibition (43). We therefore examined whether POLβ is similarly required for the maintenance/recovery of transcription following MMS treatment. To do this, we employed EdU pulse-labelling to measure global transcription and RNA polymerase I (RNAPI) immunostaining (using anti-RPA194 antibody) to measure nucleolar transcription, as described previously (43). Whereas global transcription was suppressed by ∼70% in wild type cells during a 3-hr treatment with MMS, it was suppressed by >90% in both *XRCC1^−/−^* and *POL*β *^−/−^* cells (Fig.5). Importantly, this decline in transcriptional capacity was prevented by incubation with the PARP inhibitor KU-0058948, confirming that it resulted from excessive PARP activity (Fig.5). Similar results were observed if we measured transcription in nucleolar transcription (Fig.5). Whereas RNAPI immunofoci were detected in wild type cells throughout the incubation with MMS, these foci declined by ∼90% in *XRCC1^−/−^* and *POL*β*^−/−^* cells during MMS treatment, and in both cases this decline was prevented by incubation with PARP inhibitor (Fig.5). In summary, this data suggests that like XRCC1, *POL*β is requited to prevent PARP1-dependent transcriptional suppression, during MMS treatment.

**Figure 5.**
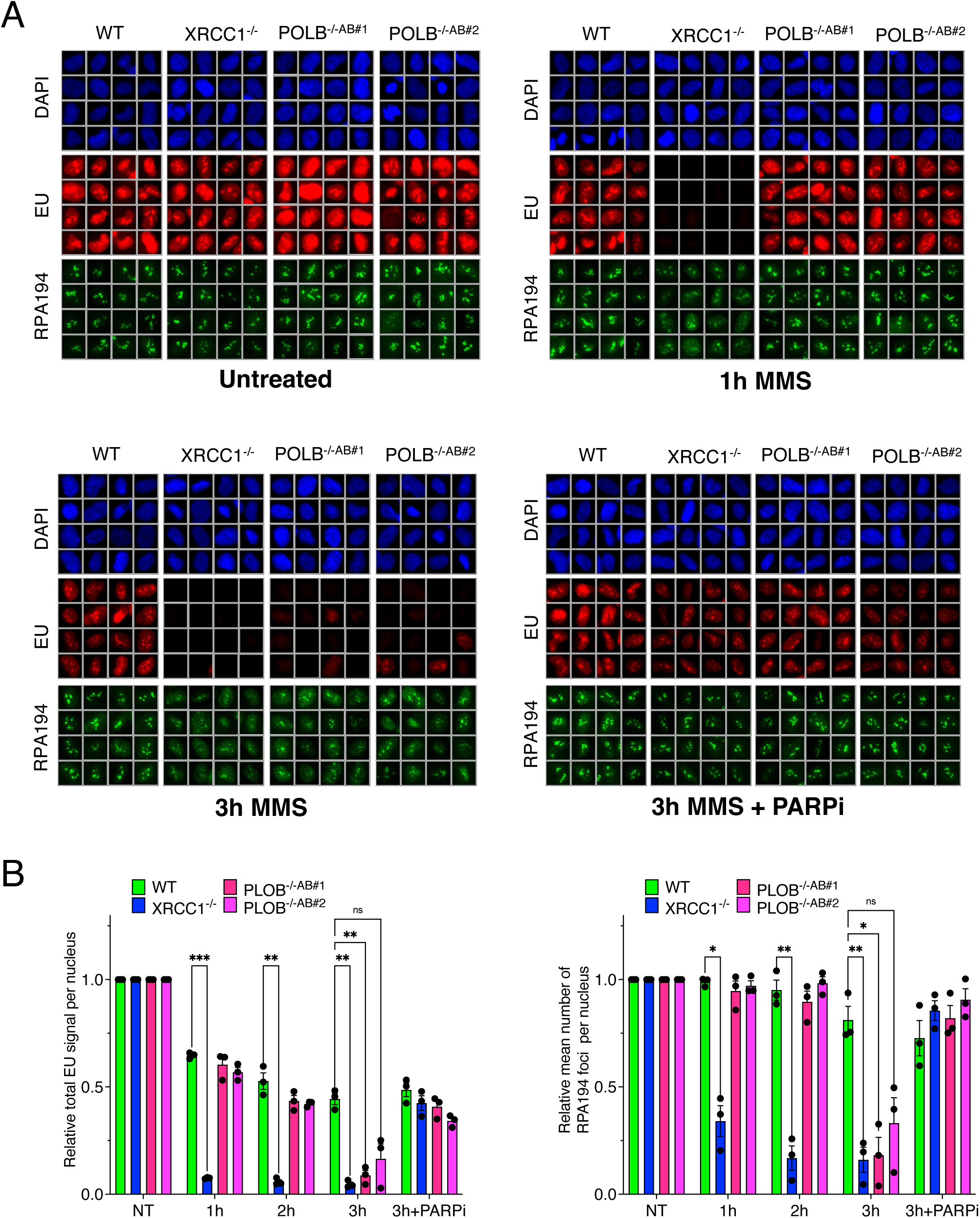
PARP-dependent suppression of transcription during BER in the absence of XRCC1 or POLβ. **(A)**, Representative galleries of scanR high content images showing the level of global transcription (EU pulse labelling) and nucleolar transcription (RNAPI/RPA194 immunostaining) in WT, *XRCC1^−/−^* (clone #3), and *POL*β *^−/−^* (clones #AB1 & #AB2) RPE-1 cells following mock treatment or at the indicated times after treatment with 0.1 mg/ml MMS. Where indicated, cells were incubated with 10 μM PARP inhibitor (PARPi; KU0058948, Axon Medchem) during MMS treatment. Cells were pulse labelled with EU for the final 20 min, prior to fixation. **(B)**, quantitation by scanR high content imaging of levels of EU pulse labelling and RPA194 immunostaining in the above experiments.

## Discussion

We showed previously that the essential role of XRCC1 in accelerating DNA base excision repair (BER) is to prevent the excessive engagement of PARP1 with BER intermediates, which otherwise impedes their repair by other BER enzymes and can lead to excessive NAD^+^ depletion and persistent PARP1 ‘trapping’ (39). This role of XRCC1 is illustrated by the greatly elevated DNA strand break intermediates that accumulate in XRCC1-defective cells during incubation with MMS, which are prevented by PARP1 deletion (39). Thus, in the absence of PARP1, XRCC1 is dispensable for rapid rates of BER. In the current work, we show that the elevated accumulation of DNA strand breaks in *POL*β *^−/−^* cells, during BER, is similarly prevented by PARP1 deletion. Thus, similar to XRCC1, the essential non-redundant role of POLβ during BER is to suppress the excessive engagement and activity of PARP1. This conclusion is consistent with previous work showing that Parp1 depletion/deletion prevents neuronal apoptosis and restores normal MMS sensitivity in *Pol*β *^−/−^* mice, although BER efficiency was not measured in these studies and it was concluded that this rescue represents the prevention of Parp1/p53-dependent apoptosis (47, 48).

It is important to note that while excessive PARP1 engagement impedes BER, some PARP activity is required for this process. This was illustrated by the increased DNA strand break intermediates that accumulated in RPE-1 cells lacking both PARP1 and PARP2, when compared to wild type RPE-1 cells. This requirement for PARP activity was independent of its role in recruiting XRCC1 or POLβ, however, because these two proteins were dispensable for rapid rates of BER in cells lacking PARP1. Rather, the requirement for PARP activity for rapid rates of BER in cells likely reflects one or more other reported role/s for PARP activity during BER, such as chromatin remodelling (49–51).

It is unclear how POLβ suppresses excessive PARP1 engagement, but it likely includes binding and/or removal of the 5’-deoxyribose phosphate from incised abasic site intermediates by the dRPase activity. This would be consistent with the reportedly high affinity of PARP1 for the incised abasic sites that are substrates for this activity, and with the observation that this dRPase activity is important for cellular resistance to MMS (21, 23, 52). Neither the dRPase activity or the DNA gap filling activities of POLβ are sufficient for suppressing PARP1 engagement, however, because this suppression also requires XRCC1. That XRCC1 and POLβ function together to prevent excessive PARP1 engagement during BER is supported by the lack of additional impact on BER rates if both XRCC1 and POLβ are deleted, when compared to XRCC1 deletion alone. In addition, recombinant XRCC1^F67A^ that cannot bind POLβ was unable to fully support rapid rates of BER. The partial rescue of BER by recombinant XRCC1^F67A^ in human *XRCC1^−/−^* cells is consistent with previous work in Chinese hamster ovary cells (44, 45), and likely reflects that XRCC1^F67A^ retains poly (ADP-ribose) binding ability and can thereby partially suppress PARP1 hyperactivity, directly, (42, 43). This may also explain why XRCC1 has a greater impact on PARP1 suppression than POLβ, because it does so by both POLβ-dependent and -independent mechanisms.

The suppression of excessive PARP1 engagement by POLβ and XRCC1 was important not only for rapid rates of BER, but also to ensure transcriptional competence following MMS-induced DNA base damage. The impact of excessive PARP1 engagement on rates of BER likely reflects the physical hindrance to access of SSB intermediates by other BER enzymes. In contrast, the impact of excessive PARP1 engagement on transcriptional competence reflects the excessive poly(ADP-ribose) polymer that is synthesised by PARP1 hyperactivity, which in turn leads to aberrant recruitment of the histone deubiquitylase and transcriptional regulator, USP3 (43). This is illustrated by the respective impact of PARP inhibitors on these two phenotypes. PARP inhibitors phenocopy the impact of XRCC1 or POLβ loss on BER rate, because they further promote the “trapping” of PARP1 on BER intermediates. In contrast, PARP inhibitors alleviate the impact of XRCC1 or POLβ loss on MMS-induced transcriptional suppression, because they prevent excessive poly (ADP-ribose) synthesis.

Collectively, our data support a model in which XRCC1 and POLβ together suppress the excessive engagement of PARP1 during BER (Fig. 6). This role is important to reduce the impact of PARP1 on the recruitment and/or engagement of other BER proteins, which otherwise cannot access BER intermediates to facilitate their repair, including XRCC1 and POLβ themselves. The mechanism by which XRCC1 and POLβ together suppress excessive POLβ activity involves their physical interaction, consistent with the idea that the rapid and efficient “hand-off” of incised BER intermediates to POLβ is important to prevent unwanted interference by PARP1. While PARP activity recruits XRCC1 protein complexes, which can then rapidly facilitate gap processing, gap filling, and DNA ligation, these enzymatic roles can be replaced by other cellular enzymes. Rather, the essential role of XRCC1 and POLβ during BER is to regulate PARP1 engagement and activity, ensuring that it does not become an obstacle to the access and repair of SSB intermediates during this important DNA repair process.

**Figure 6.**
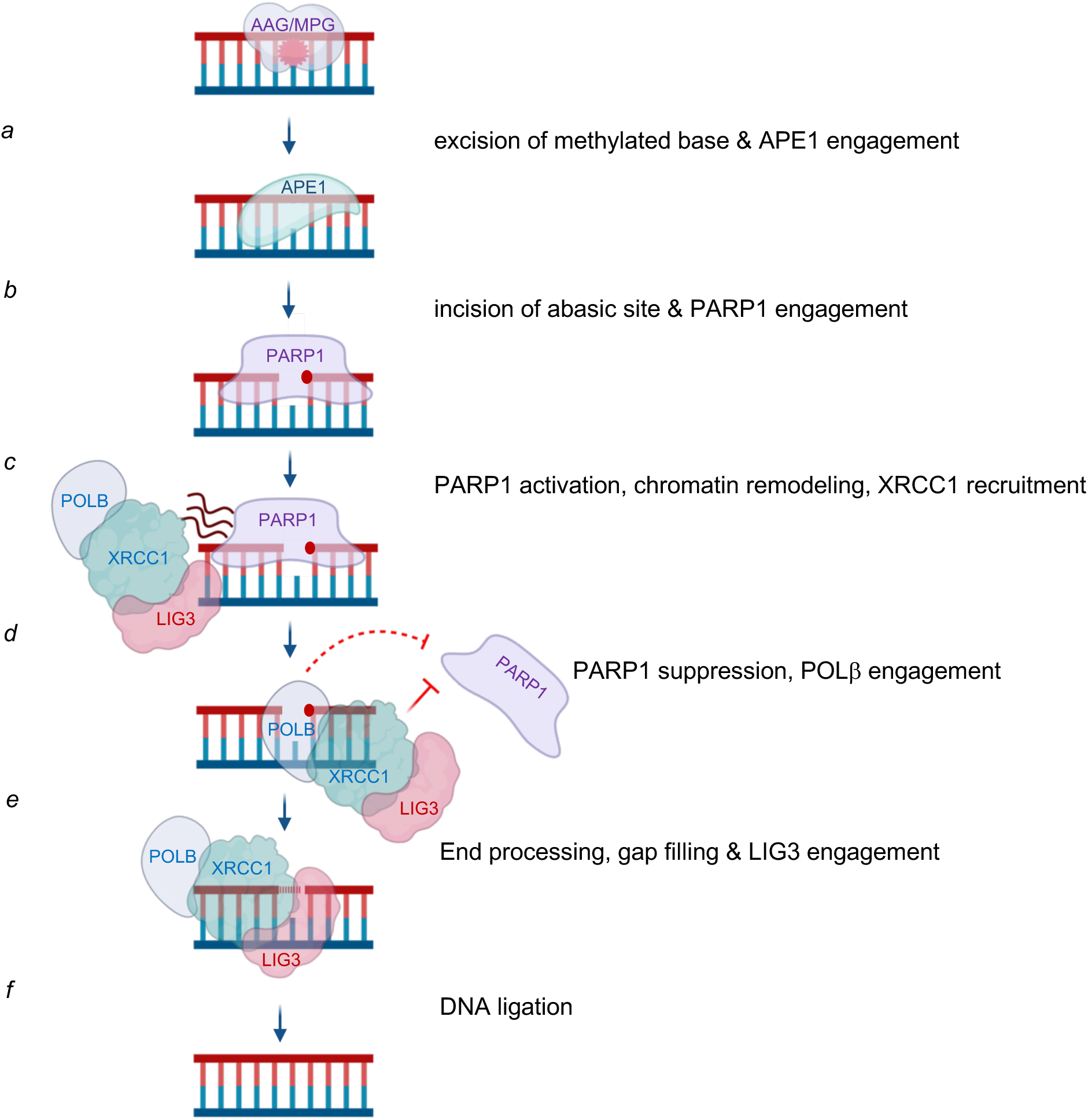
Model for XRCC1 and POLβ-dependent suppression of excessive Parp1 engagement during BER. *(a)* MMS-induced methylated bases are excised by alkyladenine/methylpurine DNA glycosylase (MPG/AAG). *(b)* The resulting abasic site is incised by AP endonuclease (APE1), creating a SSB intermediate with a 5’-deoxyribose phosphate terminus (red circle) that is detected by PARP1 (and/or PARP2). *(c)* PARP1 and/or PARP2 is activated by the SSB, resulting in PARP auto-modification, and likely also trans-modification of nearby histones (not shown), thereby enabling chromatin remodelling and the recruitment of XRCC1 protein complexes. *(d)* XRCC1 and POLβ together suppress engagement of PARP1, via a mechanism involving direct binding of XRCC1 to auto-modified PARP1 and via engagement of POLβ with the incised abasic site. (*e)* POLβ dRPase and gap filling activities remove the 5’-deoxyribose phosphate and fill the single nucleotide gap, respectively. *(f)* LIG3 ligates the remaining nick. Note that the essential role of POLβ (and XRCC1) during BER is to prevent excessive PARP1 engagement, which otherwise impedes access to the SSB intermediate by other repair enzymes. In the absence of PARP1, POLβ is dispensable for rapid rates of BER, likely because of catalytic redundancy with one or more other DNA polymerases.

## Methods

### Antibodies

The antibodies employed for this work were mouse anti-PARP1 monoclonal (Santa Cruz Cat#sc-8007; RRID:AB_628105), mouse anti-PARP1 monoclonal (Serotec clone A6.4.12), rabbit anti-PARP2 (Activ Motif 39743), rabbit anti-XRCC1 polyclonal (Novus; Cat#NBP1-87154; RRID:AB_11029388), rabbit anti-DNA polymerase beta polyclonal (Millipore Sigma #ABE1408), rabbit anti-DNA ligase III polyclonal sera (TL25), mouse anti-AAG/MPG monoclonal (Santa Cruz clone 3D1, sc-101237), anti-poly-ADP-ribose binding reagent (Millipore; Cat# MABE1031; RRID:AB_2665467), mouse anti-alpha-Tubulin monoclonal (Sigma-Aldrich; Cat#T6074; RRID:AB_477582), and rabbit anti-histone H3 polyclonal (Abcam; Cat#ab1791; RRID:AB_302613).

### Gene Edited Cells and Cell Culture

Human RPE-1 cells were incubated in a low oxygen (3%) incubator (37°C, 5% CO_2_) and maintained in DMEM-F12 Glutamax 10% FBS supplemented with penicillin/streptomycin. The gene edited RPE-1 cell lines *PARP1^−/−^* (clone #G7), *PARP2^−/−^* (clone #A1)*, PARP1^−/−^/PARP2^−/−^*(clone #E6)*, XRCC1^−/−^* (clones #3 and #23), and *PARP1^−/−^*/*XRCC1^−/−^* have been described previously (26, 53, 54). *POL*β *^−/−^* gene-edited RPE-1 cells (clones #AB1 & #AB2) were generated by the co-transfection of a Cas9-encoding and guide-encoding plasmids as described previously (53). For POLβ deletion, we co-transfected two guide plasmids simultaneously with the targeted sequences being #A (GGCCGCCATGAGCAAACGGA<u>AGG)</u> and #B (GCAGCGGGTCGTCTTCCGTG<u>GGG</u>), with PAM sequences underlined. *XRCC1*^−/−^/*POL*β ^−/−^ (clones #1 & #13), *PARP1^−/−^*/*POL*β *^−/−^* (clones #1 & #3), and *PARP1^−/−^*/*PARP2^−/−^/POL*β *^−/−^* (clone #8) double- or triple-mutant cell lines were generated by gene editing POLβ, as described above, in the *XRCC1^−/−^* (clone #3), *PARP1^−/−^* (clone #G7), and *PARP1^−/−^*/*PARP2^−/−^*(clone #E6) RPE-1 cell lines, respectively.

*MPG* ^−/−^ (clone #19) and *MPG^−/−^* /*XRCC1^−/−^* (clones #1, #3, & #9) RPE-1 cells were generated by co-transfecting of Cas9 protein and pairs of synthetic tracrRNA, using a NEON electroporator. The targeted DNA sequences in MPG were GAAATGATCACGTCTGACGA<u>TGG</u> (gRNA#1), TAGATGCTGCGGTATGGGCCCGG (gRNA#2), and GTAATGTGAACATAGCAGCG<u>AGG</u> (gRNA#3), with PAM sequences underlined. *MPG* ^−/−^ clone #19 was generated by co-transfecting gRNA#1 & gRNA#3, *MPG ^−/−^* /*XRCC1^−/−^* clone #1 by co-transfecting gRNA#1 & gRNA#2, and *MPG ^−/−^* /*XRCC1^−/−^* clones #3 & #9 by co-transfecting gRNA#2 & gRNA#3.

### Metal-chelate affinity precipitation

*XRCC1^−/−^* cells harbouring empty vector pcD2E vector or pcD2EXH plasmid encoding either wild-type histidine-tagged XRCC1(XRCC1-His), or pcD2EXH^F67A^ encoding XRCC1-His^F67A^, were stably transfected into *XRCC1^−/−^* RPE-1 clone #23 employing the NEON transfection system (Invitrogen, CA) at 1500 V 20 msec using 10µl tips. Following transfection, cells were selected against G418 (500 µg/ml) and after 10 days single colonies were isolated, amplified, and validated for expression of recombinant XRCC1.

For affinity precipitation of histidine-tagged XRCC1-His, cells were lysed in RIPA buffer (10 mM Tris/Cl pH 7.5, 150 mM NaCl, 0.5 mM EDTA, 0.1% SDS, 1% Triton_X-100, 1% deoxycholate) supplemented with 1 mM CaCl_2_, 2.5 mM MgCl_2_, 1 mg/ml DNAse (NEB), protease inhibitor cocktail (Roche), 1:1,000 benzonase (Sigma), 1:1,000 micrococcal nuclease (NEB) for 45 min on ice, and protein lysate concentrations were determined using Bradford reagent (Sigma).The lysates were then incubated for 1 hr with gentle agitation with Dynabeads™ His-Tag Isolation and Pulldown beads (20 µl Invitrogen Cat#10103D), the Dynabeads™ were premixed prior to use for 10 min in 200 ml PBS at room temperature and rinsed twice with PBS+0.02% tween 20. The Dynabeads were then washed three times with PBS+0.02% tween before incubation with 1X Laemmli buffer at 70°C for 10 min.

### siRNA and transfection

Non-targeting siRNA (ON-TARGETplus) and SMARTpool siRNA (25 nM) against PARP1 were reverse-transfected into cells using Lipofectamine® RNAiMAX (Invitrogen) according to the manufacturer’s instructions. Experiments were carried out 72 hr post-transfection.

### Chromatin fractionation & immunoblotting

Chromatin fractionation of lysed cell extracts was conducted essentially as described(39). In brief, RPE-1 cells were lysed in lysis buffer containing 100 mM or 150 mM KCl, respectively, and 50 mM HEPES pH 7.4, 2.5 mM MgCl2, 5 mM EDTA pH 8, 3 mM dithiothreitol (DTT), 0.5% Triton X-100, 10% glycerol, and protease inhibitor cocktail (Roche) for 45 min on ice. Soluble and chromatin-bound proteins were separated by centrifugation (15 min, 16,000 g), and the pellet of detergent insoluble material (including chromatin) was washed twice in lysis buffer and then sonicated to shear DNA. The soluble and chromatin extracts were mixed with SDS-PAGE sample buffer and heated for 10 min at 95°C. For protein levels in whole cell extracts (WCE), cells were lysed directly in SDS-PAGE sample buffer and heated as above. All protein extracts were subjected to SDS-PAGE followed by protein transfer onto nitrocellulose membrane and western blotting with the indicated antibodies.

To detect chromatin-bound XRCC1 by immunofluorescence, cells were rinsed in ice-cold PBS and pre-extracted in ice-cold PBS containing 0.2% Triton-X100 and 0.3 mg/ml RNAseA, for 2 min. Cells were then rinsed in PBS, fixed in 4% formaldehyde, washed in PBS (3X), and incubated in PBS containing in 0.2% Triton-X100 on ice, for a further 5 min. Cells were then blocked in PBS + 5% BSA for 1h and then incubated in PBS + 5% BSA and rabbit anti-XRCC1 polyclonal antibody (NOVUS BIO; Cat#NBP1-87154) at 1:300 dilution, for 1h. After washing with PBS (3 X 5 min), cells were incubated with an appropriate anti-rabbit secondary antibody (1:1000 in 5% BSA) for 1h, washed in PBS (3 X 5 min), counterstained with DAPI, rinsed in water to remove salt, and mounted with Mowiol solution. Images were acquired using an automated scanR imaging station.

### Immunoflourescence and transcription assays

Immunofluorescence and transcription assays were conducted essentially as described(43). In brief, cells were fixed in 4% PFA in PBS for 10 min, permeabilized in 0.2% Triton X-100 solution in PBS for 10 min, rinsed in PBS and blocked in 5% BSA solution. The cells were incubated with the appropriate primary antibody for 1 h, washed in PBS, incubated with the appropriate fluorophore-conjugated secondary antibody (Invitrogen) for 1 h and then counterstained with DAPI. To measure the global sites of nascent RNA synthesis, the cells were pulse labelled in medium containing 1 mM EU (Abcam) for 20 min before fixation. The EU pulse-labelled cells were subjected to click chemistry using a Click-iT kit (Invitrogen) according to the manufacturer’s protocol. Click chemistry was performed after the blocking step of the immunofluorescence protocol described above. Images for quantitation were acquired using the automated wide-field microscope scanR image acquisition and analysis software, with >500 cells scored per sample (Olympus). Where displayed, scanR cell galleries were generated by the Olympus Analysis Software as a representative example of the cell populations that were quantified.

### Alkaline comet assays

Cells were treated with the indicated concentration of MMS in the presence/absence of 10 mM PARP inhibitor (KU0058948) for 15 min at 37°C in medium, and alkaline comet assays conducted essentially as described previously(55). Average tail moments (an arbitrary-unit measure of DNA strand breaks) from 50-100 cells per sample were scored blind using Comet Assay IV software (Perceptive Instruments, UK), and data plotted as the mean comet tail moment (mean± S.E.M.) of these averages, from three independent experiments.

## Acknowledgements

This work was funded by an MRC Programme Grant to KWC, (MR/W024128/1), and is dedicated to my late colleague and friend Grigory Dianov.

